# Characteristic energy scales of active fluctuations in adherent cells

**DOI:** 10.1101/2022.08.17.504239

**Authors:** Avraham Moriel, Haguy Wolfenson, Eran Bouchbinder

## Abstract

Cell-matrix and cell-cell adhesion play important roles in a wide variety of physiological processes, from the single cell level to the large scale, multicellular organization of tissues. Cells actively apply forces to their environment, either extracellular matrix or neighboring cells, as well as sense its biophysical properties. The fluctuations associated with these active processes occur on an energy scale much larger than that of ordinary thermal equilibrium fluctuations, yet their statistical properties and characteristic scales are not fully understood. Here, we compare measurements of the energy scale of active cellular fluctuations — an effective cellular temperature — in four different biophysical settings, involving both single cell and cell aggregates experiments under various control conditions, different cell types and various biophysical observables. The results indicate that a similar energy scale of active fluctuations might characterize the same cell type in different settings, though it may vary among different cell types, being approximately 6 to 8 order of magnitude larger than the ordinary thermal energy at room temperature. These findings call for extracting the energy scale of active fluctuations over a broader range of cell types, experimental settings and biophysical observables, and for understanding the biophysical origin and significance of such cellular energy scales.

## I. Introduction

Noise and fluctuations play important roles in a wide variety of natural and man-made systems. In thermal equilibrium, thermal fluctuations are accurately described by equilibrium statistical thermodynamics and are controlled by the ambient temperature. Quenched noise, i.e. fluctuations and inhomogeneity in the system’s structure, is also of great importance. In living systems, which are of active nature absent in inanimate matter and which are intrinsically out-of-equilibrium, fluctuations are also highly relevant, yet are far less characterized and understood. In this brief note, we discuss the quantification of characteristic energy scales of active fluctuations (and in some cases, their statistical distributions) in adherent cells based on 4 different experimental settings, various biophysical observables and several cell types. Such fluctuations are important for a wide variety of physiological processes, from the single cell level to the multicellular organization of tissues, and their quantification is expected to significantly affect our fundamental understanding of these processes.

## II. Results

Here we describe and discuss measurements of characteristic energy scales of active fluctuations in adherent cells, adhering either to an extracellular matrix or to other cells, in 4 experimental settings and including 6 cell types in total. 2 of the experimental settings are our own and 2 are related to works available in the literature.

### A. Cell-scale orientational fluctuations of non-interacting cells under periodic forcing

In various physiological situations, cells are exposed to high-frequency periodic driving forces in addition to intrinsic and extrinsic noise, e.g. in vital conditions encountered in cardiovascular tissues in hemodynamic environments, in the lungs under breathing motion and in cardiac tissue under rhythmic heart beating. Numerous well-controlled laboratory experiments, in which living cells adhere to a deformable substrate experiencing periodic driving forces (see, for example, [1–10]), revealed that under such conditions cells reorient themselves to well-defined angles.

In such experiments, cells are exposed to a time-dependent strain tensor ***ε***(*t*) of the form 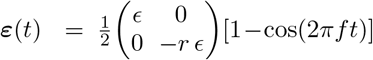, where *ϵ* is the strain amplitude in the principal direction and −*r ϵ* is the strain amplitude in the perpendicular direction and *r* is the biaxiality strain ratio, cf. Fig. 1a. The experimentalist externally controls *ϵ, r* and the periodic forcing frequency *f* (typically in the physiologically relevant range of ∼ 1 Hz). After applying the periodic forces for a long time, cells reorient themselves to an angle *θ* (or its mirror-image angle −*θ*) relative to the principal strain direction, cf. Fig. 1a. Applying this protocol to an ensemble of non-interacting cells, cf. Fig. 1a, enables the extraction of a probability distribution function *p*(*θ*) that depends on cell-scale orientational fluctuations. In Fig. 1b, we present one of the oriented cells, whose focal adhesions (green) and stress fibers (red) are colored.

**FIG. 1.**
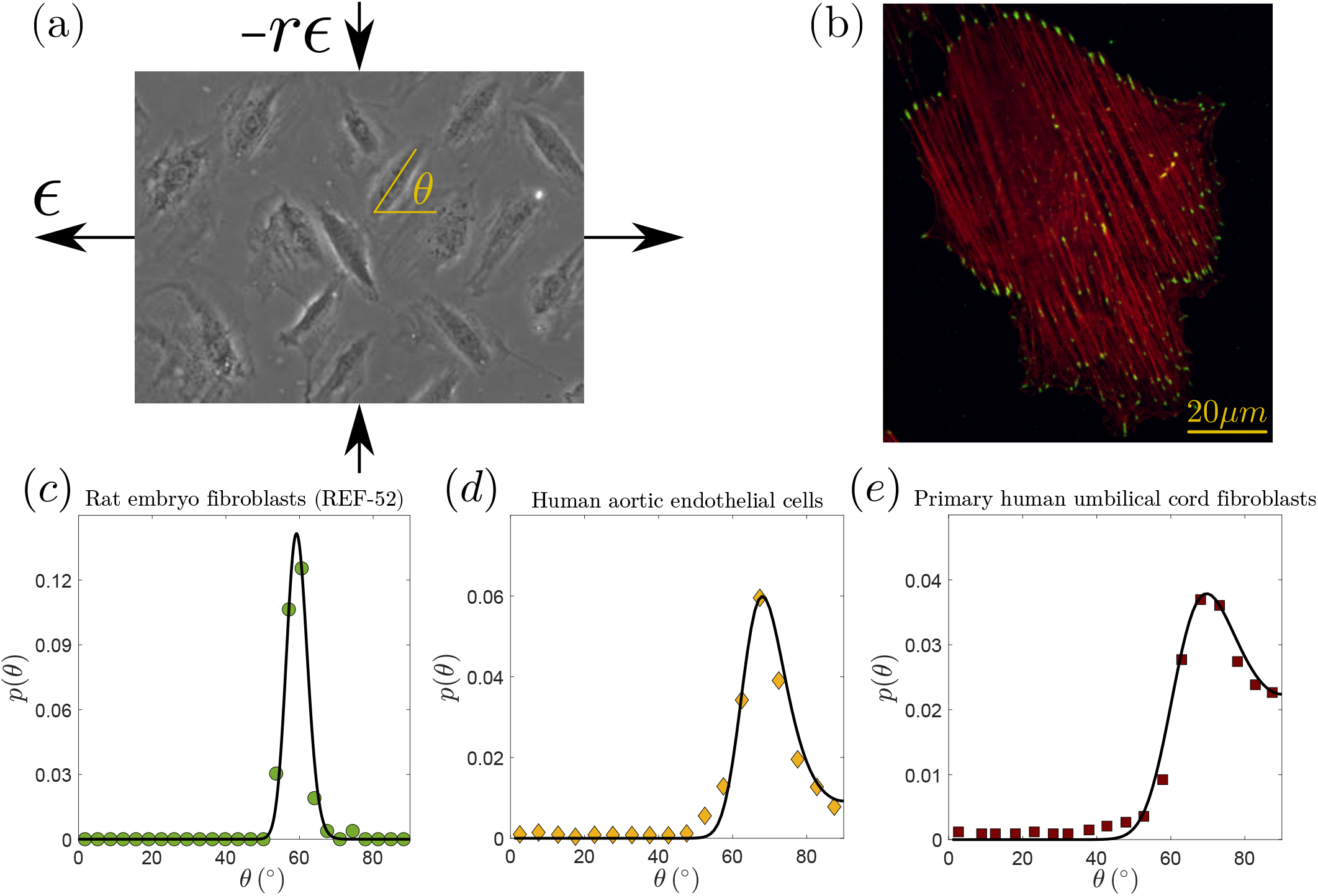
(a) A top view of REF-52 cells on a PDMS substrate under long-time periodic loading with principal strain amplitude of *ϵ*, biaxiality ratio *r* (both illustrated on the figure) and frequency *f*, see text for additional details. The cells, which initially featured random orientations, are all approximately oriented at an angle *θ* (relative to the principal strain direction, marked on the figure) or its mirror-image angle −*θ* (not marked) in the long-time limit. (b) A zoom in on a single REF-52 in its oriented state (after 6 hours of periodic stretching), showing its stress fibers (red) and focal adhesions (green) [10]. (c) Measured *p*(*θ*) for REF-52 (green circles) under long-time periodic stretching with *ϵ* = 10.4%, *r* = 0.46 and *f* = 1.2 Hz (adapted from Fig. 3b of [10]). The solid line corresponds to Eq. (1), with *b* = 1.15 extracted from the maximum of the distribution (see text for details) and a single fitting parameter set to (*k*_B_*T*_eff_)*/*(*V*_cell_ *E*_cell_) = 3.2 × 10^*−*5^. (d) Measured *p*(*θ*) for human aortic endothelial cells (orange diamonds) under periodic stretching with *ϵ* = 10.0%, *r* = 0.34 and *f* = 0.5 Hz (see Fig. 5B in [5] and Fig. 3a in [10]). The solid line corresponds to Eq. (1), with *b* = 1.23 extracted from the maximum of the distribution (see text for details) and a single fitting parameter set to (*k*_B_*T*_eff_)*/*(*V*_cell_ *E*_cell_) = 6.2 × 10^*−*5^. (e) Measured *p*(*θ*) for primary human umbilical cord fibroblasts (brown squares) under periodic stretching with *ϵ* = 31.7%, *r* = 0.28 and *f* = 9 × 10^*−*3^ Hz (see Fig. 4 in [4] and Fig. 2f in [10]). The solid line corresponds to Eq. (1), with *b* = 1.18 extracted from the maximum of the distribution (see text for details) and a single fitting parameter set to (*k*_B_*T*_eff_)*/*(*V*_cell_ *E*_cell_) = 1.6 × 10^*−*3^.

A recent theory [9–11] predicted that *p*(*θ*) takes the form

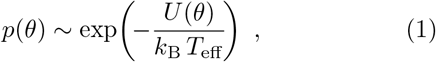

where *T*_eff_ is an effective temperature associated with active orientational fluctuations, *k*_B_ is Boltzman’s constant and *U* (*θ*) is an orientation-dependent cell-scale elastic energy. The latter takes the form [9, 10] 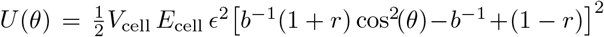, where *V*_cell_ is the volume of the cell and *E*_cell_ is its coarsegrained elastic modulus. *b* is a dimensionless number that quantifies the elastic anisotropy of the cell [9, 10], related to the most probable angle (i.e. the maximum of *p*(*θ*) attained at *θ*_m_) through 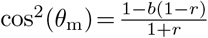 [9, 10].

The probability density *p*(*θ*) is theoretically predicted up to the unknown dimensionless combination (*k*_B_ *T*_eff_)*/*(*V*_cell_ *E*_cell_) [10], which can be then extracted from experiments. This procedure has been pursued very recently in [10], and we report some examples here. In Fig. 1c, we present *p*(*θ*) for rat embryo fibroblasts (REF-52, the experimental data correspond to Fig. 3b in [10]). The solid line corresponds to the theoretical prediction in Eq. (1) obtained with a single parameter fit, yielding (*k*_B_ *T*_eff_)*/*(*V*_cell_ *E*_cell_) = 3.2 × 10^*−*5^. In order to transform the latter into an effective temperature in physical units, we need estimates of *V*_cell_ and *E*_cell_. The area of the REF-52 shown in Fig. 1b is roughly 3000 *µ*m^2^ and its height is about 2 *µ*m, leading to *V*_cell_ ≈ 6 × 10^*−*15^ m^3^ (note that *V*_cell_ is likely to reveal cell-cell variability and follow its own statistical distribution, which is not taken into account here). The coarse-grained elastic modulus of REF-52 under these biophysical conditions was estimated to be *E*_cell_ ≈ 10 kPa [10]. Taken together, we obtain *T*_eff_ ∼ 10^8^ K, which is about 6 orders of magnitude larger than room temperature.

Our next goal is to see whether and to what extent *T*_eff_ might differ for different cell types under approximately the same biophysical conditions. In Fig. 1d, we present *p*(*θ*) for human aortic endothelial cells under similar periodic driving conditions (the experimental data correspond to Fig. 3a in [10], which were extracted from Fig. 5B in [5]). The solid line corresponds to the theoretical prediction in Eq. (1) obtained with a single parameter fit (see the values of the other, known parameters in the figure caption), yielding (*k*_B_ *T*_eff_)*/*(*V*_cell_ *E*_cell_) = 6.2 × 10^*−*5^, a similar dimensionless value as the one obtained for REF-52. Assuming that *V*_cell_ and *E*_cell_ are roughly similar to the corresponding values of REF-52, we conclude that the two cell types feature roughly the same *T*_eff_ under similar biophysical conditions.

To further test the degree of generality of this interesting result, we present in Fig. 1e *p*(*θ*) for primary human umbilical cord fibroblasts under similar periodic driving conditions (the experimental data correspond to Fig. 2f in [10], which were extracted from Fig. 4 in [4]). The solid line corresponds to the theoretical prediction in Eq. (1) obtained with a single parameter fit (see the values of the known parameters in the figure caption), yielding (*k*_B_ *T*_eff_)*/*(*V*_cell_ *E*_cell_) = 1.6 × 10^*−*3^. Assuming again that *V*_cell_ and *E*_cell_ are roughly similar to the corresponding values of the two other cell types considered above, one obtains *T*_eff_ ∼ 10^10^ K, which is about 2 orders of magnitude larger than the value estimated for the two other cell types considered above (and about 8 orders of magnitude larger than room temperature). Interestingly, it has been recently shown that primary human umbilical cord fibroblasts feature a dramatically different intrinsic timescale (related to reorientation dynamics, not discussed here) compared to REF-52 [10]. It would be interesting to explore in future work whether this difference is related to the enhanced level of fluctuations they exhibit. One way or the other, the results indicate that *T*_eff_ might be cell type dependent and is always many orders of magnitude larger than room temperature.

**FIG. 2.**
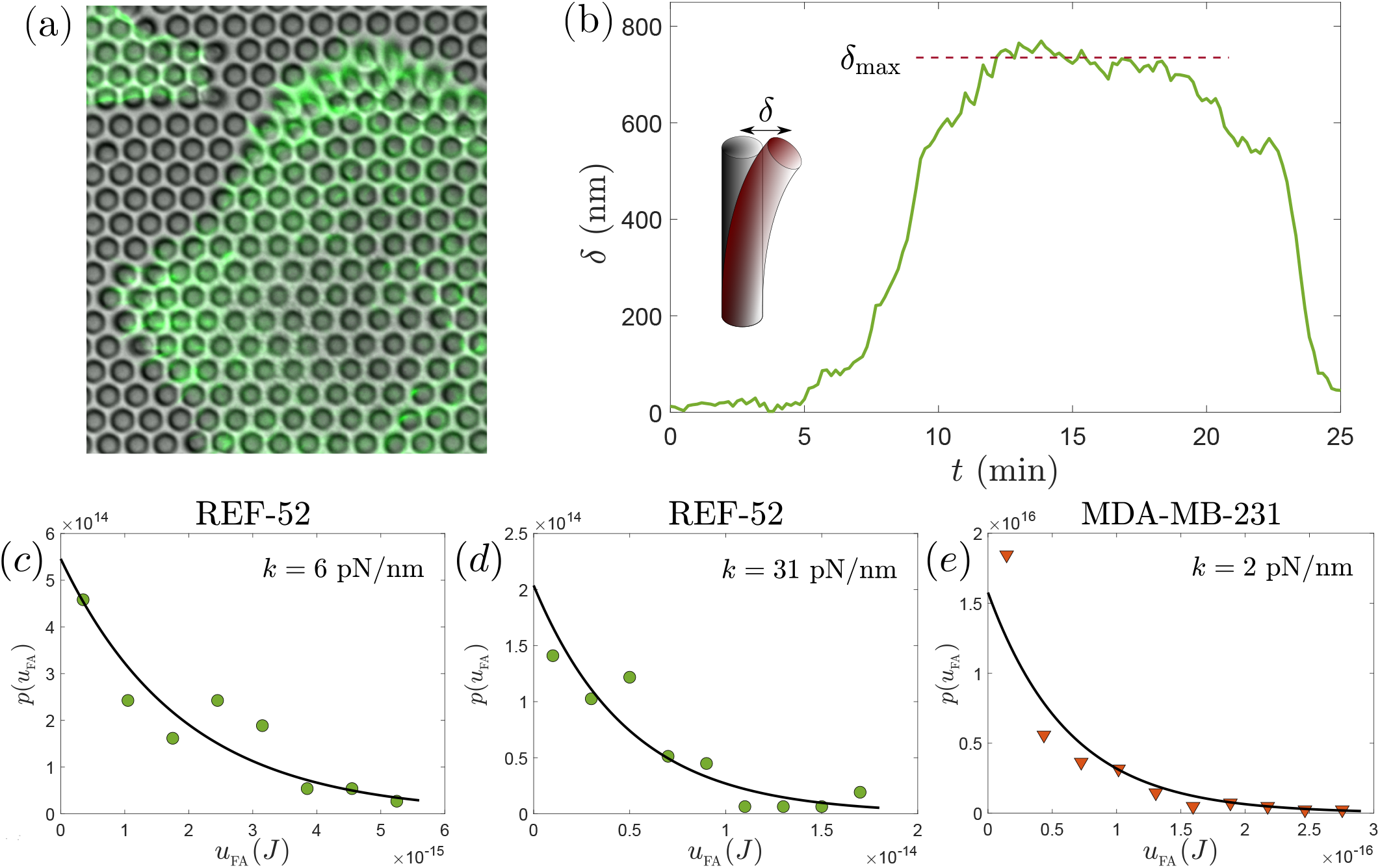
(a) A top view of a REF-52 cell, whose F-actin is shown in green, adhering to a micropillar array (the micropillar’s diameter is 2 *µ*m and the pillars center-to-center distance is 4 *µ*m) in its steady spreading state (a portion of another REF-52 is observed at the top-left corner). (b) An example of a displacement *δ* of a micropillar (here of bending rigidity *k* = 31 pN/nm) as a function of time *t* in the steady spreading state. The displacement (deflection) *δ* is illustrated in the inset (the magnitude of *δ* is exaggerated for visual clarify). Here *t* = 0 is chosen such that the buildup of force/displacement as a focal adhesion starts to assemble around *t* = 5 min is observed. *δ*(*t*) reaches a maximal displacement *δ*_max_ (marked on the figure), where it plateaus for ∼ 10 min., before the focal adhesion starts to disassemble, until the micropillar is released around *t* = 25 min. (c) Measured *p*(*u*_FA_) for REF-52 (green circles), where 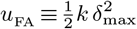, with *k* = 6 pN/nm. The solid line corresponds to Eq. (2) with a single fitting parameter set to *T*_eff_ = 1.4 × 10^8^ K. (d) The same as panel (c), but for *k* = 31 pN/nm. The solid line corresponds to Eq. (2) with a single fitting parameter set to *T*_eff_ = 3.6 × 10^8^ K. (e) Measured *p*(*u*_FA_) for human breast adenocarcinoma cells (MDA-MB-231, orange triangles), with *k* = 6 pN/nm. The solid line corresponds to Eq. (2) with a single fitting parameter set to *T*_eff_ = 4.5 × 10^6^ K.

### B. Adhesion-scale contractile energy fluctuations of stationary adherent cells

Our next goal is to test whether the same cell type features the same *T*_eff_ under different biophysical settings and observables under consideration. To that aim, we consider REF-52 cells adhering to an array of fibronectin-coated, flexible, PDMS (polydimethylsiloxane) micropillars [12], cf. Fig. 2a, without applying external driving forces. The micropillars resist deflection according to the Euler-Bernoulli beam theory, featuring bending rigidity *k* = 3*πER*^4^*H*^*−*3^*/*4, where *R* is the pillar’s radius, *H* is the pillar’s height and *E* is the Young’s modulus of PDMS (see the caption of Fig. 2 for the values of *R* and *k*, as well as for the micropillar center-to-center distance).

The cells are allowed to adhere to the micropillar array for a sufficiently long period of time, until they reach a steady spreading state. The latter steady state, where the cell features a constant area on average, is still subjected to fluctuations. In particular, in the steady spreading state, focal adhesions are formed at micropillars and apply to them a contractile force, resulting in a displacement of magnitude *δ*(*t*) as a function of time *t*. As shown in Fig. 2b, *δ*(*t*) initially rises and reaches a characteristic maximal value *δ*_max_, and after leveling off for a characteristic time of 10 minutes it decreases until the focal adhesion disassembles altogether and reassembles at a nearby micropillar (the latter is not shown).

We then define a characteristic adhesion-scale contractile energy as 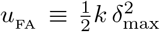, where the subscript FA stands for focal adhesion. It is important to note that *u*_FA_ is a contractile energy defined at the single focal adhesion scale (i.e. it characterizes the energy the cell invests in deflecting a single pillar through a single focal adhesion) and generated by the cell, as opposed to *U*(*θ*) of Eq. (1), which is a cell-scale elastic energy related to the external driving force.

Considering an ensemble of non-interacting cells in the steady spreading state, i.e. accounting for both intra-cell and inter-cell variations, we construct the probability distribution function *p*(*u*_FA_), presented in Fig. 2c for a micropillar array with *k* = 6 pN/nm. The distribution approximately follows a Boltzmann-like form

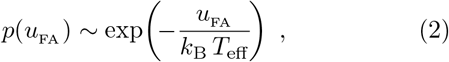

which readily allows to extract *T*_eff_ in its natural physical units. The latter is a clear advantage as it does not involve the uncertainties associated with *V*_cell_ and *E*_cell_, which were required in the previous subsection in order to transform a dimensionless effective temperature to a dimensional one. A Boltzmann-like fit of Eq. (2) with *T*_eff_ = 1.4 × 0^8^ K is added to Fig. 2c in the solid line. Quite remarkably, *T*_eff_ = 1.4 × 10^8^ K coincides with the value of *T*_eff_ extracted in Fig. 1c for the same cell type, but for a different biophysical setting and a different probed observable. This result suggests that a similar energy scale of fluctuations may characterize the same cell type in different situations, a possibility that clearly calls for additional investigation.

To further look into this issue, we present in Fig. 1d *p*(*u*_FA_) for REF-52, in exactly the same experimental setting as in Fig. 1c, but for a 5-fold larger micropillar bending rigidity, i.e. *k* = 31 pN/nm. It is crucial to stress that the substrate rigidity is varied by geometry — *k* in this experimental paradigm is varied by varying the micropillar height *H* alone, keeping its radius *R*, the substrate modulus *E* (here of PDMS) and the surface chemistry (here the fibronectin coating) fixed; this is one of the greatest advantages of the micropillar array paradigm [12]. The distribution is again Boltzmann-like, and fit to Eq. (2) with *T*_eff_ = 3.6 × 10^8^ K is added to Fig. 2d in the solid line. While this *T*_eff_ is not 5-fold different from that of the *k* = 6 pN/nm value, it clearly indicates that the energy scale of fluctuations is not an entirely intrinsic property.

All of the cell types considered above form rather strong adhesions and apply significant forces to the extracellular matrix — through stress fibers — or to each other — through cell-cell bonds. It would be then interesting to consider yet another cell type that is known to attach weakly to the microenvironment and to posses few stress fibers. In this case, we expect *T*_eff_ to be significantly smaller. To that aim, we consider human breast adenocarcinoma cells MDA-MB-231 (an epithelial, human breast cancer cell line), which is a model cell line for invasive cancer. These cells are very motile, and possess very few stress fibers, if at all (see, for example, [13]). We apply the same experimental paradigm to MDA-MB-231, using a micropillar array with *k* = 2 pN/nm. The results are presented in Fig. 2e, where a Boltzmann-like fit of Eq. (2) with *T*_eff_ = 4.5 × 10^6^ K is added in the solid line. This *T*_eff_ value is ∼ 2 orders of magnitude smaller than the values obtained for REF-52 under similar conditions, in agreement with our expectation.

### C. Energy fluctuations of a single cell moving inside an aggregate of other cells

Next, we briefly consider a different set of experiments available in the literature, where *T*_eff_ is also estimated. In [14], the motion of individual pigmented retinal epithelial cells in aggregates of neural retinal cells (obtained from chicken embryos) has been tracked. The individual cells undergo a random walk, characterized by a linear diffusion coefficient *D* ≃ 3 × 10^*−*12^ cm^2^/sec = 3 × 10^*−*16^ m^2^/sec [14]. The viscosity of the aggregate was measured to be *η* ≃ 10^5^ Pa·sec [15]. Then, assuming an equilibrium-like Stokes-Einstein relation for the linear diffusion of one liquid inside another of the form *T*_eff_ = 5*π D η a/*(2 *k*_B_), yielded an estimate of *T*_eff_ ≃ 1.4 × 10^8^ K using *a* ≃ 8 *µ*m for the linear dimension of the cell [15]. Very interestingly, this energy scale of fluctuations roughly coincides with the *T*_eff_ obtained for REF-52 and human aortic endothelial cells in very different biophysical situations.

### D. Interfacial energy fluctuations during cell sorting in a mixture of two cell-type aggregates

Experiments related to those briefly reported in the previous subsection studied cell sorting in a mixture of the same two types of cells (i.e. pigmented retinal epithelial cells and neural retinal cells [15]). Employing an analogy between cell sorting and the separation of immiscible fluids allowed to extract an energy fluctuations scale characterizing the interface between two cell aggregates.

For two immiscible fluids at thermal equilibrium, the ordinary temperature *T* is related to the interfacial tension *σ*_t_ through 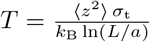 in two dimensions. Replacing *T* by *T*_eff_, and using the measured mean-squared-displacement ⟨*z*^2^⟩ ≃ 0.04*a*^2^ perpendicular to the interface (i.e. ⟨*z*^2^⟩ is a measure of interfacial tissue fluctuations, where ⟨·⟩ is an average over the entire interfacial area), the measured neural retina-pigmented epithelium interfacial tension *σ*_t_ ≃ 10 dyne/cm = 10^*−*2^ N/m, the measured linear size of the pigmented epithelial cell aggregate *L* ≃ 200 *µ*m and the measured linear dimension of the cell *a* ≃ 8 *µ*m (as in the previous subsection) one obtains *T*_eff_ ≃ 5.8 × 10^8^ K. This value is in the same ballpark as the one obtained in the previous subsection, which also coincides with the *T*_eff_ obtained for REF-52 and human aortic endothelial cells, yet again indicating that cells might feature the same scale of energy fluctuations in different settings.

## III. Discussion

The results presented in this brief note indicate that active cellular fluctuations of adherent cells can be systematically quantified in various biophysical settings. Such analyses can provide quantitative information about the statistical distribution of relevant biophysical observables and to estimate an effective temperature *T*_eff_ that defines a characteristic energy scale. The latter is found to be many orders of magnitude larger than the ordinary thermal energy scale, reflecting both the out-of-equilibrium, active nature of the fluctuations and the relatively large energy scales associated with cellular adhesion structures.

We find that various cell types under different biophysical conditions and observables being probed are characterized by a characteristic energy scale of fluctuations of *T*_eff_ ∼ 10^8^ K, i.e. about 6 orders of magnitude larger than room temperature. Yet, we also find that another cell type — which is known to exhibit dramatically different intrinsic temporal dynamics associated with its adhesions during cellular reorientation — features a characteristic energy scale of fluctuation of *T*_eff_ ∼ 10^10^ K. This significantly larger *T*_eff_ suggests that cell type dependence cannot be generally excluded. Furthermore, we show that when adhesion structures and contractile forces are significantly weaker, as in the cancerous cells considered in Fig. 2e, a much smaller *T*_eff_ emerges.

These findings call for extracting the energy scale of active fluctuations over a broader range of cell types, experimental settings and biophysical observables, and for understanding the biophysical origin and significance of such cellular energy scales. Once done, it may be possible to develop a better understanding of the collective spatiotemporal dynamics of adherent cells interacting with an extracellular matrix and among themselves.

## Acknowledgements

We are grateful to Sam Safran for pushing us to compare the energy scale of active fluctuations across different biophysical settings. We are grateful to Ariel Livne for stimulating discussions and for providing most useful comments on the manuscript. E.B. acknowledges support from the Ben May Center for Chemical Theory and Computation and the Harold Perlman Family. H.W. acknowledges support from the Israel Science Foundation and from the Rappaport Family Foundation. H.W. is an incumbent of the David and Inez Myers Career Advancement Chair in Life Sciences.

